# Purine and pyrimidine analogues differentially regulate cell wall precursor biosynthesis to control β-lactam susceptibility in methicillin resistant *Staphylococcus aureus*

**DOI:** 10.64898/2026.05.24.727567

**Authors:** Aaron C. Nolan, Sarah Byrne, Merve S. Zeden, James P. O’Gara

## Abstract

Maintaining the efficacy of β-lactam antibiotics against *Staphylococcus aureus* is a clinical priority given the prevalence of methicillin-resistant *S. aureus* (MRSA). We previously showed that the pyrimidine analogues 5-fluorouracil (5-FU) and 5-fluorouridine (5-FUrd) synergize with β-lactams. Here, we extended this by evaluating additional nucleotide metabolism-targeting agents. Gemcitabine (Gem) and mitomycin C (Mito), like 5-FU and 5-FUrd, exhibited intrinsic anti-MRSA activity and potentiated β-lactams, whereas the purine analogue 6-thioguanine (6-TG) showed distinct, often antagonistic effects. Transcriptomic analysis revealed that pyrimidine-targeting agents repress lysine and glutamate biosynthesis, while 6-TG induced these pathways, implicating amino acid metabolism in β-lactam potentiation. Consistent with this, pyrimidine analogues also suppressed GlmS expression, potentially limiting UDP-GlcNAc production required for cell wall synthesis, and synergized with fosfomycin. Fluorescence microscopy confirmed that the potentiation of oxacillin activity by pyrimidine-targeting agents, but not 6-TG, was accompanied by impaired peptidoglycan synthesis. Additionally, glutathione-mediated attenuation of killing implicated reactive oxygen species in the bactericidal activity of cloxacillin combinations. Finally, these agents displayed strong anti-biofilm activity, further enhanced in combination with daptomycin and rifampicin. Together, these findings highlight the potential of pyrimidine analogues to potentiate cell wall-targeting antibiotics and identify an important role for modulation of cell wall precursor pathways in this anti-MRSA activity.

**Importance:** Drug interactions can complicate the treatment of antimicrobial resistant infections in patients undergoing treatment for cancer highlighting the importance of understanding the effects of anti-cancer drugs on pathogens like MRSA. Here, we investigated several drugs that target nucleotide metabolism and are used to treat cancer, fungal, and viral infections, both alone and in combination with commonly used penicillin-type antibiotics. We found that pyrimidine analogue drugs enhanced the activity of these antibiotics against MRSA, whereas the purine analogue 6-thioguanine reduced antibiotic effectiveness. These drugs altered the bacterial cell wall and other metabolic pathways linked to antibiotic susceptibility. Our findings reveal the potential to repurpose certain anticancer drugs to improve treatment of MRSA infections, while also cautioning that some drug combinations may interfere with antibiotic therapy.

## Introduction

Ameliorating the challenge posed by multidrug-resistant pathogens (1, 2) is a major public health priority. Management of infections caused by methicillin resistant *Staphylococcus aureus* (MRSA), which remains one of the most important Gram-positive pathogens (3), typically necessitates vancomycin, daptomycin or linezolid due to their high-level resistance to β-lactams (4–6). However, resistance to these antimicrobial drugs and failure to eradicate certain infection types remain significant challenges (7–10). Biofilm and persister cells further complicate the treatment of MRSA infections, leading to increased rates of therapeutic failure (11–15). Antibiotic adjuvants that modulate nucleotide homeostasis can increase MRSA susceptibility to β-lactams and anti-folate antibiotics, attenuate virulence and eradicate biofilm (16, 17). The multifaceted interrelationship between nucleotide metabolism, antimicrobial resistance and virulence (18–24) further supports the utility of targeting these pathways with adjuvant therapeutics.

Nucleotide analogues that interfere with nucleotide metabolism are used in the treatment of cancer, viral infections and fungal infections (25–27), raising the prospect that they may also be useful in the treatment of infections caused by bacterial pathogens including MRSA (28). Consistent with this possibility, several nucleotide analogue anticancer drugs display antibacterial activity, including inhibition of peptidoglycan synthesis (29, 30). Given that patients with cancer are frequently immunocompromised and more vulnerable to infection and sepsis (31), the therapeutic potential of anti-cancer drugs with antibacterial activity may be particularly important (32).

Anti-cancer drugs which disrupt DNA synthesis can generally be categorised as antimetabolites of nucleotides/nucleosides required for DNA replication or drugs that directly bind to DNA target sequences triggering chemical changes and DNA damage (27, 33, 34). The conservation of nucleotide metabolism in eukaryotes and prokaryotes underpins the antibacterial activity of these agents (28). The commonly used anticancer drug 5-fluorouracil (5-FU) has been shown to produce thymidine related antimetabolites that interfere with DNA and RNA processing (27). For example, in MRSA, disruption of thymidine biosynthesis, which is likely to be impacted by 5-FU, affects intracellular signalling, pyrimidine homeostasis, and daptomycin susceptibility (35, 36).

In this study, we evaluated the anti-MRSA activity of a panel of anticancer drugs that act as antimetabolites or disrupt nucleotide metabolism, including 6-thioguanine (6-TG), 5-FU, 5-fluorouridine (5-FUrd), gemcitabine (Gem), and mitomycin C (Mito), as well as the antifungal 5-fluorocytosine (5-FC) and the antiviral agents acyclovir (Acy) and azidothymidine (AZT) (37–43) (Fig. 1). The potential of these drugs to potentiate antibiotics targeting the cell wall was also determined and supported by fluorescence microscopy using a cell wall dye. RNAseq analysis performed with MRSA strain JE2 exposed to 5-FU, 5-FUrd, Gem, Mito and 6-TG provided insights into their mechanism of action. Comparative genomics was used to identify mutations associated with increased resistance to these drugs. The contribution of reactive oxygen species (ROS) arising from DNA damage induced by these drugs was assessed using the ROS quencher glutathione in antibiotic time-kill assays. Finally, the antibiofilm activity of several anti-cancer drugs used alone or in combination with other antibiotics was measured. The data presented reveal the potential of these anticancer drugs as part of a novel therapeutic regimen for MRSA infections.

**Fig. 1.**
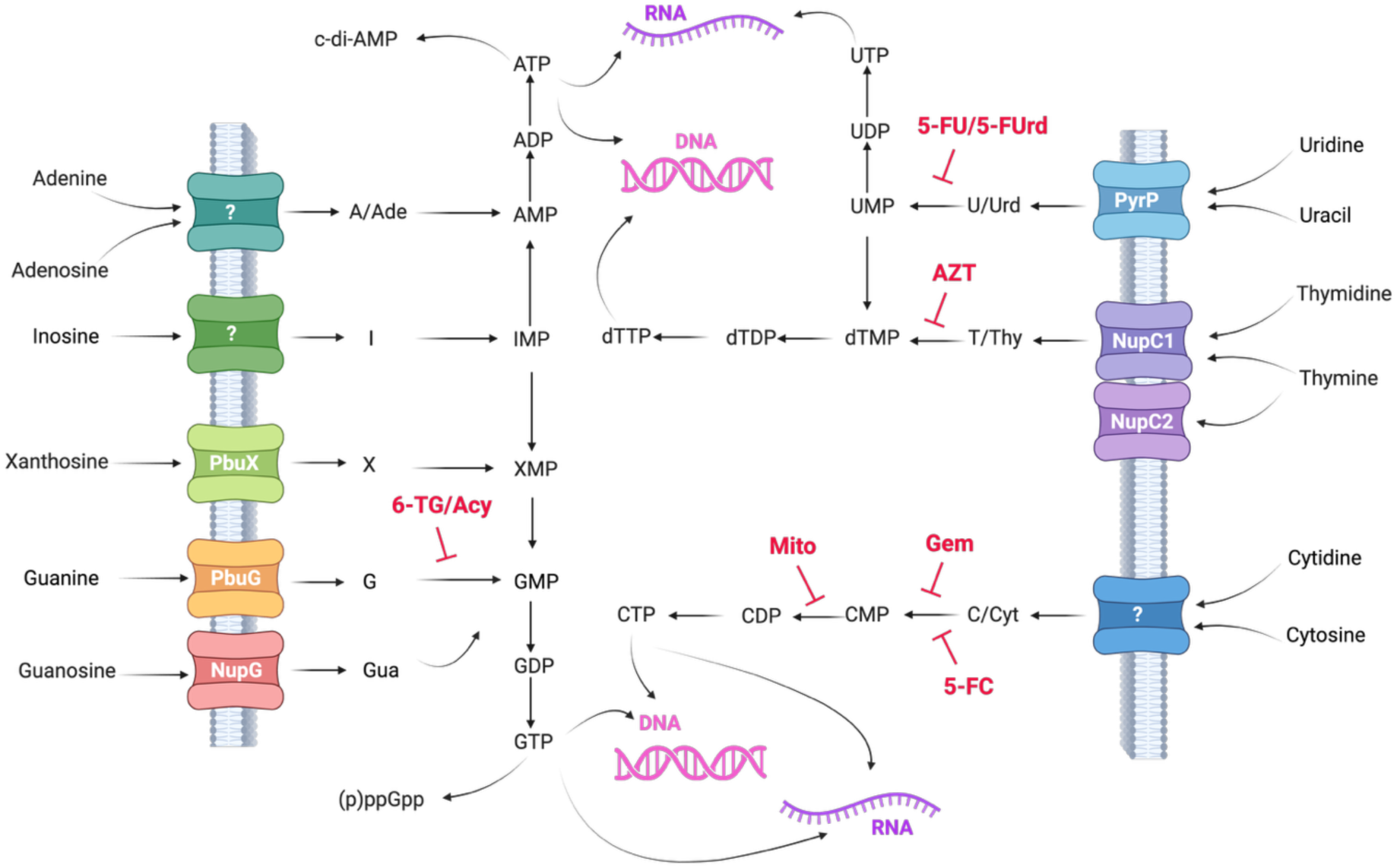
Overview of nucleotide salvage pathways in *S. aureus* responsible for producing DNA/RNA from exogenous nucleotides/nucleosides. Purines can be transported by known transporters PbuG, NupG and PbuX (16, 44), and fluxed to GTP and ATP. GTP and ATP can be converted to (p)ppGpp or c-di-AMP, respectively or used for DNA/RNA synthesis. Pyrimidine nucleotides and nucleosides are salvaged by PyrP, NupC1 and NupC2 (45) and converted into deoxy trinucleotides before being incorporated into DNA (dCTP, dTTP) or RNA (dCTP, dUTP). Predicted inhibition of enzymes involved in pyrimidine biosynthesis by 5-FC, Mito, Gem, AZT, 5-FU, 5-FUrd and 6-TG is depicted by red block arrows.

## Results

### Anti-cancer drugs targeting pyrimidine metabolism have anti-MRSA activity

We recently reported that the antibiotic adjuvant guanosine potentiates the activity of the pyrimidine antimetabolites 5-fluorouracil (5-FU) and 5-fluorouridine (5-FUrd), which are used as anti-cancer drugs (17). Here we extended these experiments to investigate the anti-MRSA activity of several drugs targeting nucleotide metabolism that are used in the treatment of viral and fungal infections, and cancer (Fig. 2). Minimum inhibitory concentration (MIC) assays revealed that MRSA strains JE2, MW2, BH1CC and COL were susceptible to 5-fluorouracil (5-FU), 5-fluorouridine (5-FUrd), 6-thioguanine (6-TG), gemcitabine (Gem) and mitomycin C (Mito), but not to acyclovir (Acy), azido-thymidine (AZT) and 5-fluorocytosine (5-FC) (Table 1). Consistent with this, growth of JE2 was significantly impaired in 0.5× MIC, 1× MIC and 2× MICs of 5-FU, 5-FUrd, 6-TG, Gem and Mito, whereas 1× MICs of 5-FC, Acy and AZT only marginally affected growth (Fig. 2A-F). Similarly, 1× MICs of 5-FU, 5-FUrd, Gem, Mito and 6-TG also significantly inhibited the growth of the MRSA strains BH1CC, MW2 and COL (Fig. S1).

**Fig. 2.**
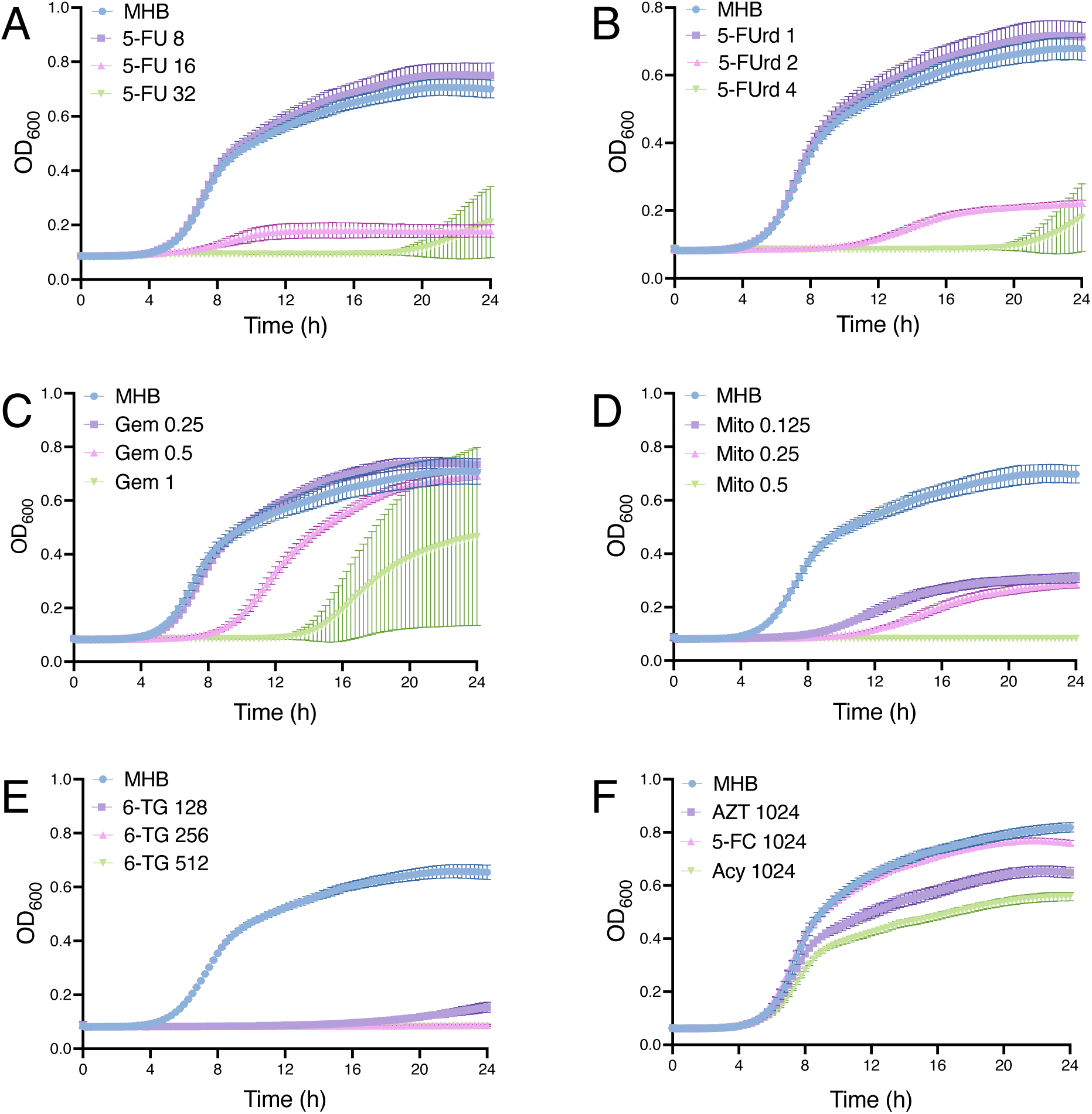
Inhibition of MRSA growth by clinically used anti-cancer compounds. **A-F.** Growth of JE2 for 24 h at 37°C in Mueller-Hinton broth (MHB) and MHB supplemented with (A) 5-fluorouracil (5-FU) 0.5× MIC (8 µg/ml), 1× MIC (16 µg/ml) or 2× MIC (32 µg/ml), (B) 5-fluorouridine (5-FUrd) 0.5× MIC (1 µg/ml), 1× MIC (2 µg/ml) or 2× MIC (4 µg/ml), (C), gemcitabine (Gem) 0.5× MIC (0.25 µg/ml), 1× MIC (0.5 µg/ml) or 2× MIC (1 µg/ml), (D), mitomycin C (Mito) 0.5× MIC (0.125 µg/ml), 1× MIC (0.25 µg/ml) or 2× MIC (0.5 µg/ml), (E) 6-thioguanine (6-TG) 0.5× MIC (128 µg/ml), 1× MIC (256 µg/ml) or 2× MIC (512 µg/ml) and (F) 1024 µg/ml azidothymidine (AZT), 1024 µg/ml 5-fluorocytosine (5-FC) or 1024 µg/ml acyclovir (Acy). Growth (OD600) was measured at 15 min intervals in a Tecan plate reader. Data are the average of 3 independent experiments plotted using GraphPad Prism V9 and error bars represent standard deviation.

**Table 1.** MRSA minimum inhibitory concentrations (µg/ml) for 5-flourouracil (5-FU), 5-flourouridine (5-FUrd), 6-thioguanine (6-TG), mitomycin C (Mito), gemcitabine (Gem), azidothymidine (AZT), acyclovir (Acy) and 5-fluorocytosine (5-FC).

| MRSA strain | Compound |  |  |  |  |  |  |  |
| --- | --- | --- | --- | --- | --- | --- | --- | --- |
|  | 5-FU | 5-FUrd | 6-TG | Mito | Gem | Acy | AZT | 5-FC |
| JE2 | 16-32 | 1-2 | 128-256 | 0.25 | 0.5 | >1024 | >1024 | >1024 |
| MW2 | 32 | 4 | 512 | 0.03 | 1 | >1024 | >1024 | >1024 |
| BH1CC | 16 | 2 | 256 | 0.03-0.06 | 0.5-1 | >1024 | >1024 | >1024 |
| COL | 4-8 | 1-2 | 256 | 0.008-0.015 | 0.5-1 | >1024 | >1024 | >1024 |

To investigate compensatory mutations underlying the emergence of increased resistance to 5-FU, 5-FUrd, and Gem after 12-20 h (Fig. 2A-C), JE2 was passaged in MHB containing 0.5× MIC of each compound, and whole-genome sequencing was performed on 2-3 independent resistant mutants per drug (Table S1). Consistent with previous work (44), mutations in *hpt* (purine salvage) were identified in two 6-TG resistant mutants. A 5-FU-resistant mutant carried a Thr_12_Ile substitution in a predicted Na⁺-dependent transporter, suggesting that it may play a role in uracil transport, while others harboured mutations in a transcriptional regulator or a glutathione S-transferase. 5-FUrd resistance involved either large (16-gene) deletions or a SNP upstream of the *ald1* operon. Gem resistance mapped to pyrimidine metabolism genes (*cmk*, *nupC*, *dck*) and upstream of *nrdI*. Mito resistance was associated with mutations in *rpoC*, as well as *pdhB*, *agrC*, *ftsK*, and *phoP*. These findings highlight the potential of anticancer drugs for MRSA treatment, notwithstanding the capacity for resistance to emerge.

### Exposure of MRSA to drugs targeting pyrimidine metabolism is associated with repression of amino acid biosynthesis genes involved in cell wall metabolism

Exposure of MRSA to drugs targeting pyrimidine metabolism represses amino acid biosynthetic pathways linked to cell wall metabolism. RNAseq analysis revealed striking differences in transcriptional responses of MRSA treated with 0.5× MICs of 5-FU, 5-FUrd, Gem and Mito compared to the purine analogue 6-TG (Fig. S2 and Fig. 3A-B). In general, pyrimidine-targeting drugs elicited transcriptional responses opposite to those induced by 6-TG. The pyrimidine analogues 5-FU, 5-FUrd and Gem produced broadly similar cell wall expression profiles, whereas Mito showed a more distinct pattern (Fig. 3A). Overall, 294, 142, 388, 133, and 293 genes were differentially expressed following exposure to 5-FU, 5-FUrd, 6-TG, Gem and Mito, respectively (Fig. S2).

**Fig. 3.**
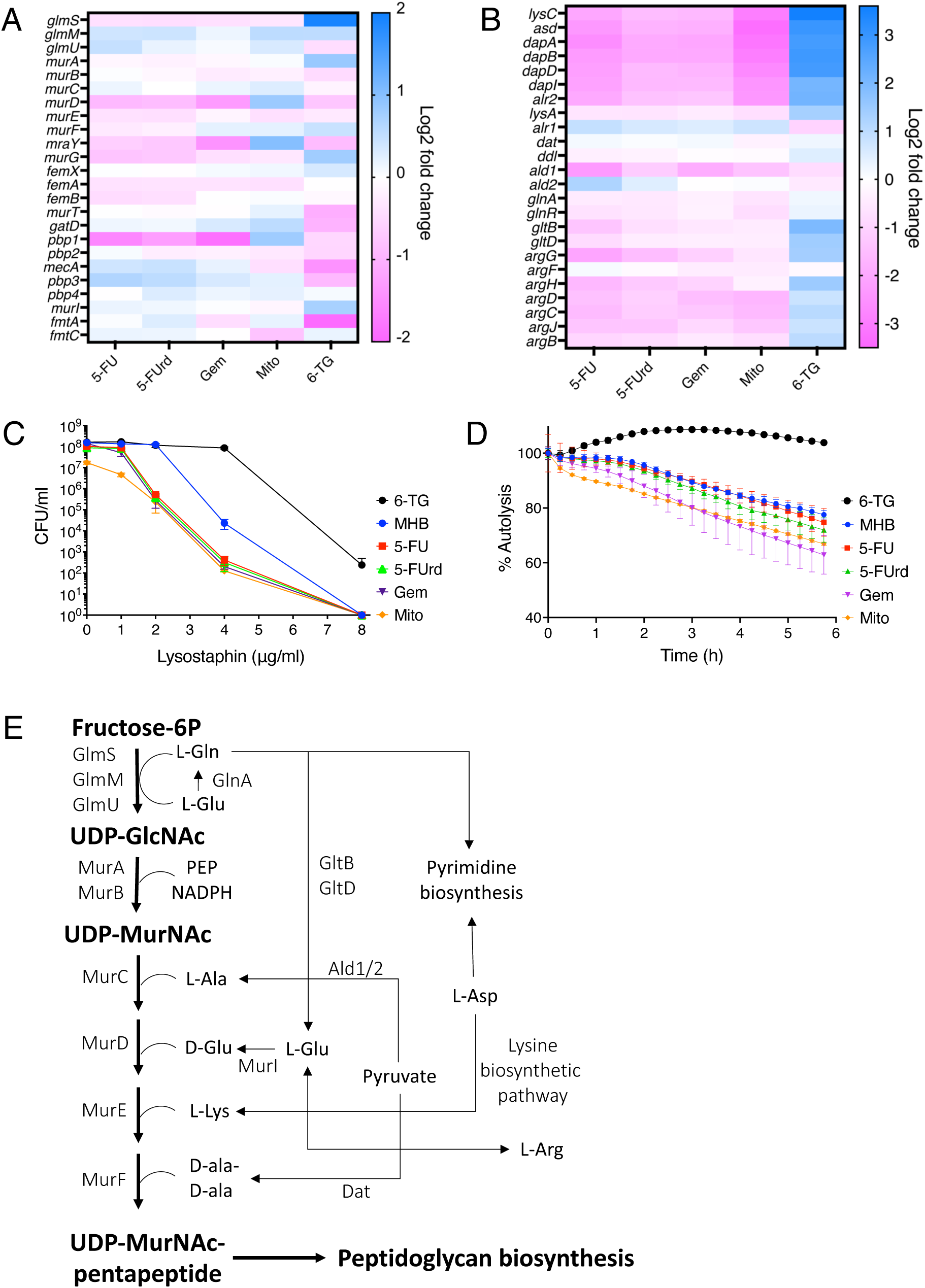
Drugs targeting pyrimidine metabolism repress transcription of genes impacting peptidoglycan biosynthesis. **A-B.** Heat map analysis of (A) cell wall synthesis and (B) amino acid biosynthesis gene expression following 1 h exposure to 5-FU, 5-FUrd, 6-TG, Gem and Mito compared to untreated control. Three independent RNASeq experiments were conducted, and average data were plotted using GraphPad Prism V9. **C.** Lysostaphin time-kill assay. Exponential phase JE2 cells that had been grown in MHB alone or MHB supplemented with 0.25× MICs of 5-FU (4 µg/ml), 5-Furd (0.25 µg/ml), Gem (0.125 µg/ml), Mito (0.0625 µg/ml), or 6-TG (32 µg/ml) were adjusted to a starting cell density of approximately 1×10^8^ CFU/ml in fresh TSM supplemented to lysostaphin 0, 1, 2, 4 and 8 µg/ml, and CFUs enumerated after 90 min. The average data from three independent experiments with standard deviations plotted using GraphPad Prism V9 are presented. **D.** Triton X-100-induced autolysis of JE2 grown in MHB alone or MHB supplemented with 0.25× MICs of 5-FU (4 µg/ml), 5-Furd (0.25 µg/ml), Gem (0.125 µg/ml), Mito (0.0625 µg/ml), or 6-TG (32 µg/ml) to an OD_600_ of 0.5 in MHB at 37°C before being washed in cold PBS and resuspended in 0.1% Triton X-100 with the OD_600_ adjusted to 1. The OD_600_ was monitored at 15-min intervals, and autolysis was expressed as a percentage of the initial OD_600_. The average data from three independent experiments with standard deviations plotted using GraphPad Prism V9 are presented.(**E.** Intersection of peptidoglycan, pyrimidine and amino acid biosynthetic pathways regulated by pyrimidine anti-metabolite drugs.

Among cell wall-associated genes, *glmS*, which catalyses the first step in peptidoglycan biosynthesis, was downregulated by pyrimidine-targeting agents but upregulated by 6-TG (Fig. 3A). Consistent with this, genes involved in arginine biosynthesis, metabolically linked to glutamine and glutamate metabolism, were similarly repressed by pyrimidine-targeting drugs and induced by 6-TG (Fig. 3B). The *glnRA* operon followed the same pattern (Fig. 3B). Disruption of glutamine synthesis has previously been shown to increase β-lactam susceptibility in MRSA via reduced amidation of iso-D-glutamate in the peptidoglycan stem peptide (46, 47).

Genes involved in the biosynthesis of lysine (*lysA* operon) and glutamate (*gltBD*), which are essential components of the peptidoglycan stem peptide were also repressed by 5-FU, 5-FUrd, Gem and Mito, but upregulated by 6-TG (Fig. 3B). Upregulation of the alanine racemase gene *alr1*, required for D-alanine production, by pyrimidine-targeting drugs was not consistent with the broader repression of cell wall-associated pathways, whereas 6-TG exposure again produced the opposite effect (Fig. 3B). Interestingly, differential modulation of lysine and glutamate gene expression by the anti-cancer drugs was inversely associated with lysostaphin susceptibility (Fig. 3C), suggesting that the availability of these amino acids influences the structure and/or accessibility of the pentaglycine cross-bridges in peptidoglycan that are the target of lysostaphin (48). Furthermore, 6-TG, but not the drugs targeting pyrimidine metabolism, repressed autolytic activity in JE2 (Fig. 3D), serving as further evidence of altered peptidoglycan structure. Together, these data indicate that pyrimidine-targeting agents impair peptidoglycan precursor biosynthesis through repression of key amino acid pathways (Fig. 3E), providing a mechanistic insight into their potentiation of cell wall-targeting antibiotics.

### Drugs targeting pyrimidine metabolism potentiate the anti-MRSA activity of β-lactam antibiotics and fosfomycin

Consistent with RNAseq data indicating modulation of peptidoglycan biosynthetic pathways, 5-FU, 5-FUrd, Gem, and Mito exhibited synergy with the β-lactams oxacillin (OX) and cloxacillin (CLOX) (Fig. 4A-D). In contrast, the purine analogue 6-TG antagonised the activity of both OX and CLOX (Fig. 4E). Synergy was also observed between fosfomycin, which inhibits MurA, and 5-FU (FIC = 0.375), 5-FUrd (FIC = 0.375), and Mito (FIC = 0.365), whereas 6-TG was antagonistic (FIC = 2). Killing assays further revealed that growth of JE2 in sub-MIC 5-FU, 5-FUrd, Gem or Mito, alone or in combination with sub-MIC OX, enhanced fosfomycin killing whereas growth in 6-TG or OX/6-TG had had no effect on fosfomycin killing (Fig. S3).

**Fig. 4.**
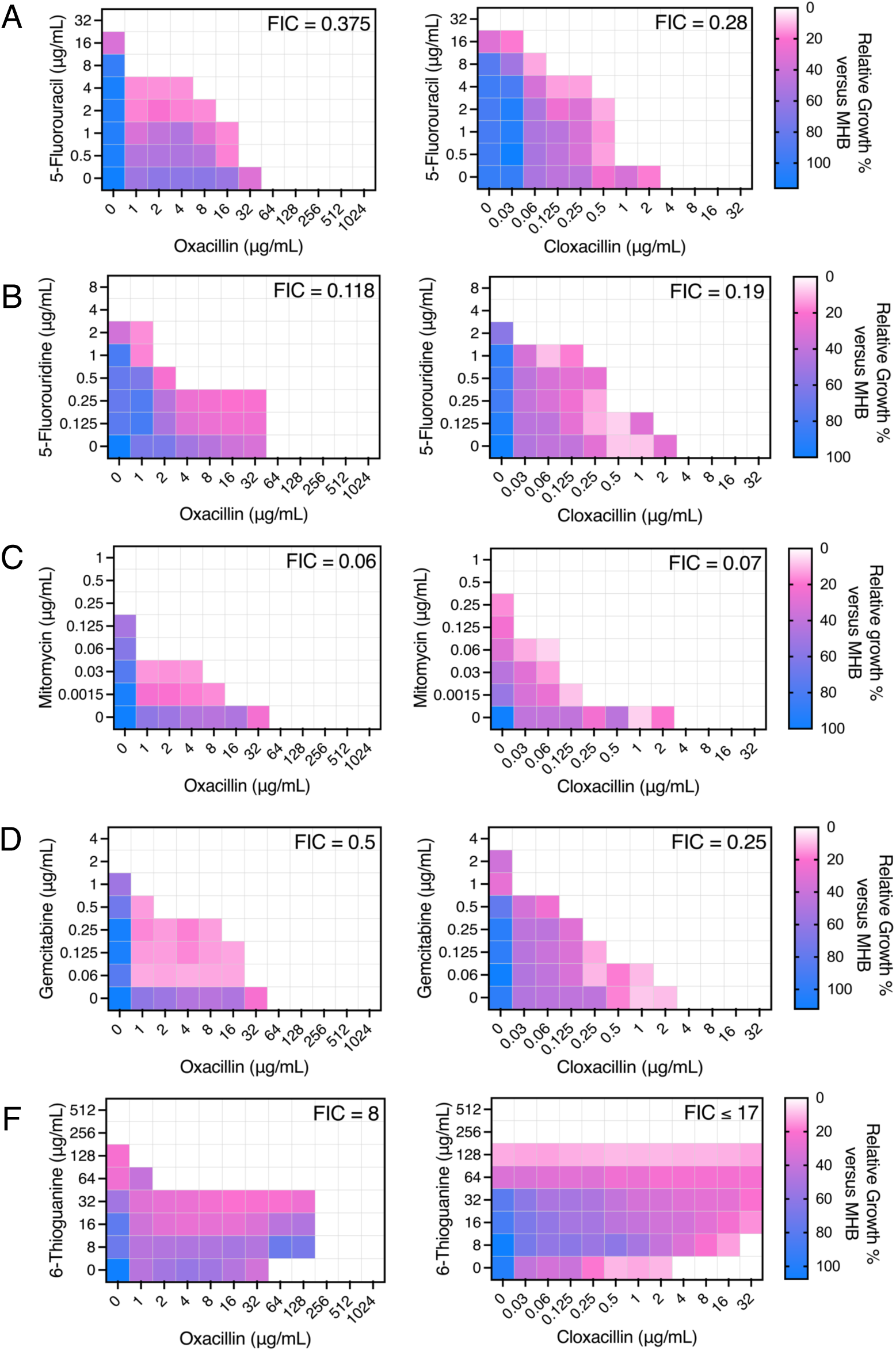
5-FU, 5-FUrd, Gem and Mito (but not 6-TG) potentiate the activity of oxacillin (OX) and cloxacillin (CLOX) against MRSA. **A-E.** Checkerboard titration assays were conducted to measure drug synergy between (A) 5-FU, (B) 5-FUrd, (C) Gem, (D) Mito and (E) 6-TG and the β-lactam antibiotics OX and CLOX against JE2 grown for 24 h in MHB in 96-well plates. Relative growth in the presence of the antibiotics is shown by heat map compared to growth in MHB (100% growth). Experiments were repeated three times and representative plates are shown. FIC values ≤0.5 are considered synergistic, values between 0.5-2 are indifferent and values >2 are antagonistic. Heatmaps were generated using GraphPad Prism V9.

Fluorescence microscopy with the D-alanine analogue HADA, which is incorporated into the cell wall, and the nucleic acid stain SYTO 9 was used to visualise the morphological impact of exposing MRSA strain JE2 to 0.125× MICs of the anti-cancer drugs alone and in combination with OX for 1 h. HADA-labelled JE2 exposed to 5-FU, 5-FUrd, Gem and Mito was similar to MHB control indicative of no significant effect on cell wall biosynthesis (Fig. 5A,B). In contrast, cells treated with 6-TG alone exhibited reduced HADA labelling indicative of slower cell wall turnover (Fig. 5B), perhaps reflective of reduced autolysis (Fig. 3D). Cells exposed to Mito had visibly reduced SYTO 9 staining, which is consistent with the predicted DNA damage and reduced DNA synthesis (Fig. 5C)(49). As shown previously, exposure to OX alone was associated with increased cell size. HADA fluorescence was lower in OX-treated cells than MHB grown cells indicative of altered HADA incorporation and SYTO 9 staining of OX-treated cells was similar to control cells grown in MHB. The synergy between OX and 5-FU, 5-FUrd, Gem and Mito was associated with reduced HADA labelling in JE2 cells treated with these combinations compared to cells grown in comparison to lone compounds. In contrast, consistent with the capacity of 6-TG to antagonise OX activity, HADA and SYTO 9 straining of JE2 cells treated with the OX/6-TG combination was similar to cells grown in 6-TG alone (Fig. 5A,C). Analysis of cell division plane and septation phenotypes revealed that, similar to OX, 5-FU, 5-FUrd, Gem, and Mito increased the proportion of MRSA cells with aberrant septal morphology and division defects during growth, whereas 6-TG-treated cells exhibited few detectable abnormalities relative to MHB-grown controls (Fig. 5D). Combination treatment with OX and 5-FU, 5-FUrd, Gem, or Mito further increased the proportion of cells exhibiting aberrant septation and division phenotypes, whereas OX/6-TG-treated cells displayed few detectable septation defects (Fig. 5D). These data support the hypothesis that pyrimidine-targeting agents enhance antibiotic efficacy by disrupting peptidoglycan biosynthesis, whereas 6-TG antagonises OX-mediated interference with this pathway.

**Fig. 5.**
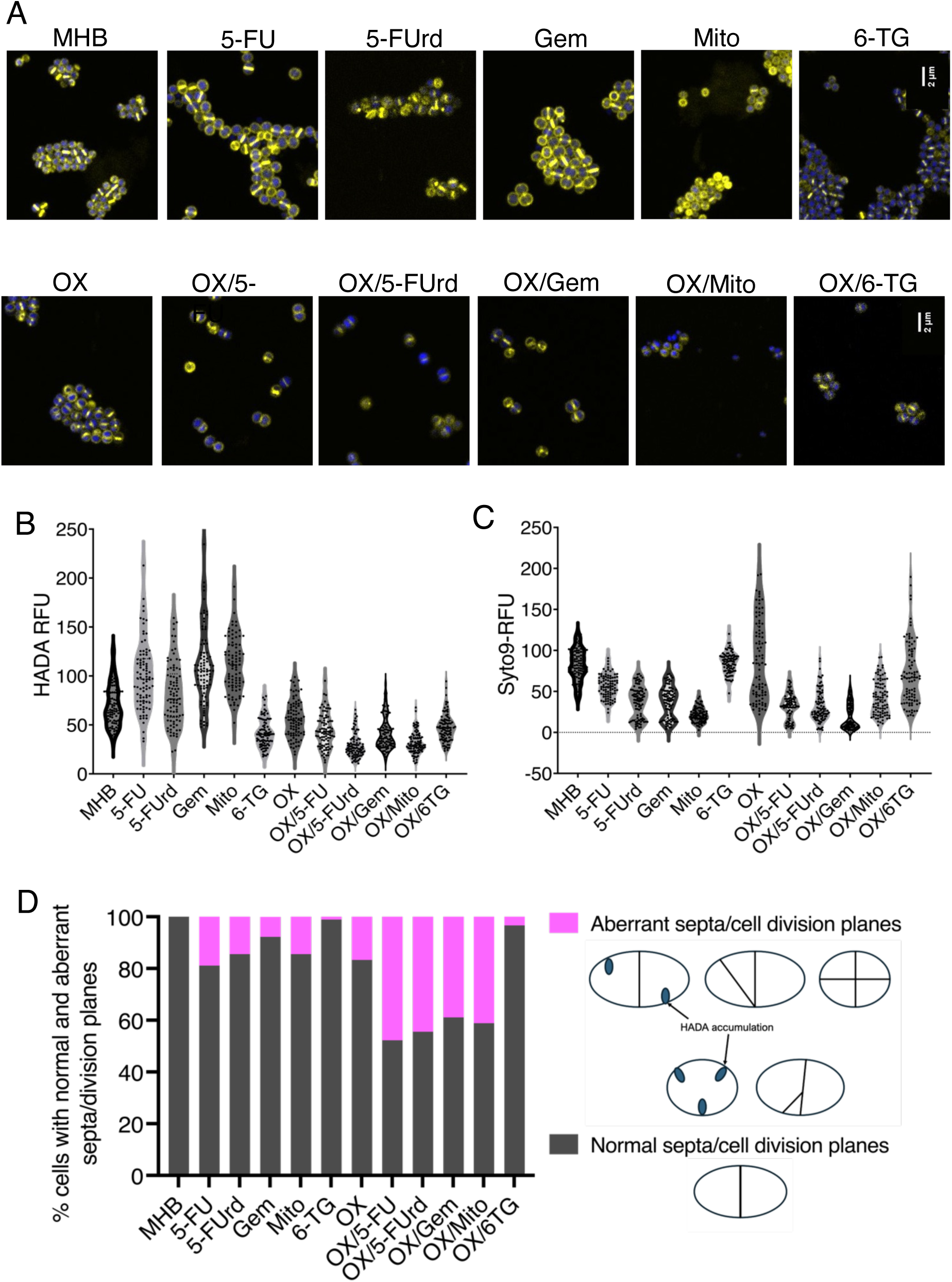
Combinations of OX with 5-FU, 5-FUrd, Gem and Mito but not 6-TG interfere with cell wall biosynthesis. **A.** Representative microscopic images captured after 1 h exposure of JE2 to anti-cancer drugs alone (**top**) and in combination with OX (**bottom**). JE2 cells were exposed to the drugs at 1× MICs before being labelled with HADA (yellow) to label D-alanine incorporation into peptidoglycan and SYTO 9 (blue) to label DNA. Images of JE2 cells from three biological replicates were acquired and representative images are shown. **B-C.** Violin plots quantifying (B) HADA and (C) SYTO 9 fluorescence intensity, expressed as relative fluorescence units (RFU), in JE2 cells treated with 5-FU, 5-FUrd, Gem, Mito and 6-TG alone and in combination with OX. HADA is a fluorescent D-alanine analogue incorporated into newly synthesised peptidoglycan and SYTO9 is a nucleic acid stain. **D.** Quantification of the relative proportion of cells exhibiting normal or aberrant septation phenotypes following treatment with 5-FU, 5-FUrd, Gem, Mito and 6-TG alone and in combination with OX. Representative schematics illustrate the major morphological phenotypes observed, including abnormal septal placement, altered division planes, and incomplete septa. Data are representative of 90 cells analysed from at least 3 independent experiments (30 per biological replicate).

### The reactive oxygen species (ROS) scavenger glutathione partially reverses the bactericidal activity of cloxacillin/anticancer drug combinations

Consistent with the predicted effects of Mito and other anti-cancer drugs on DNA integrity, several genes involved in DNA damage response and repair were differentially regulated (Fig. 6A). Specifically, 5-FU, 5-FUrd, Gem, and Mito downregulated *sosA*, *uvrA*, *uvrB*, and *recA*, while *recN* was upregulated (Fig. 6A). SosA functions as a cell division inhibitor activated in response to DNA damage and is proposed to interact with key cell division proteins, including PBP1 (penicillin-binding protein 1), DivIC, and PBP3 (52). In contrast, 6-TG did not significantly alter the expression of any of these genes (Fig. 6A).

**Fig 6.**
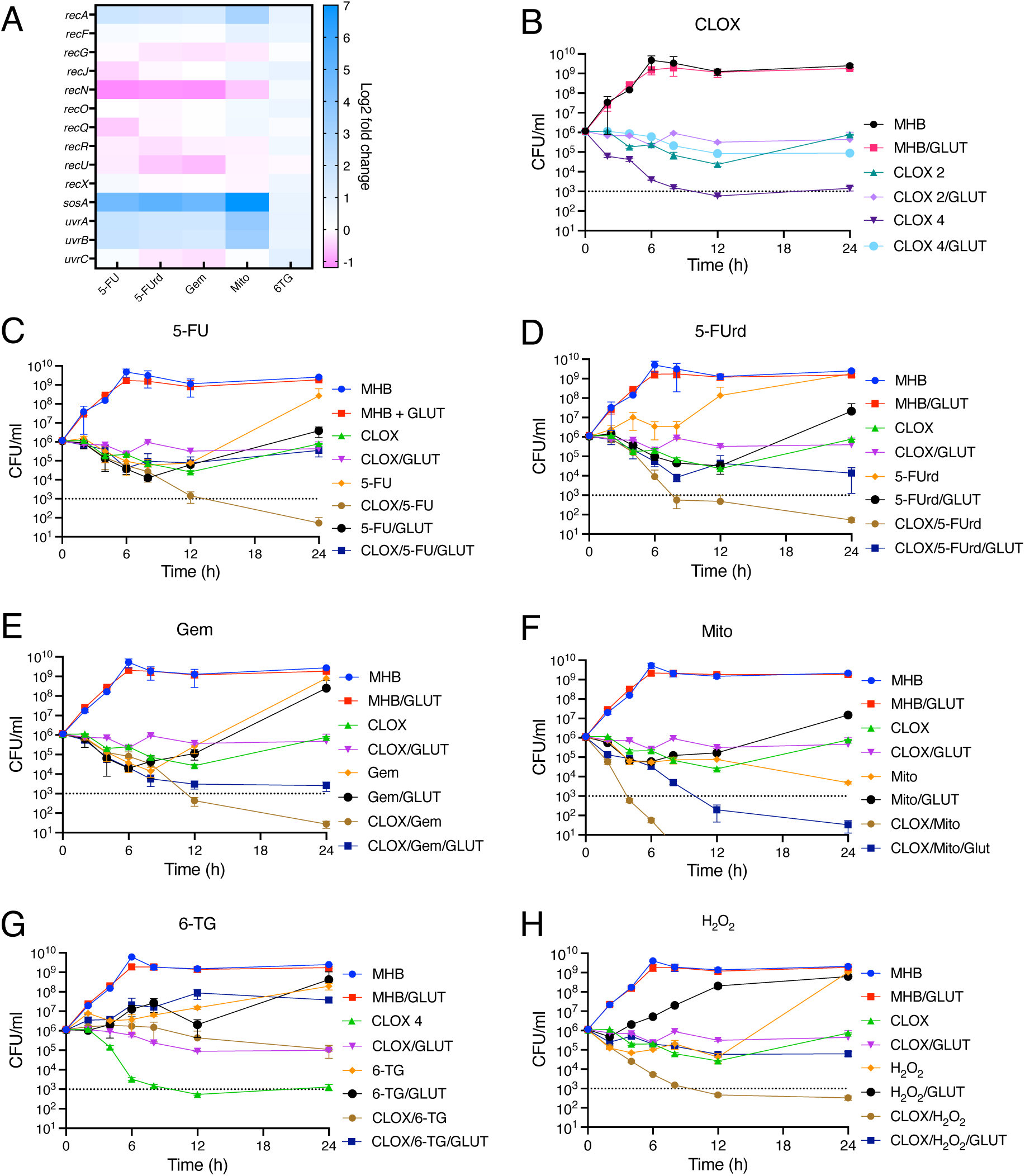
Glutathione partially reverses the bactericidal activity of cloxacillin/anticancer drug combinations against MRSA. **A.** Heat map analysis of DNA damage response gene expression following 1 h exposure to 5-FU, 5-FUrd, Gem, Mito and 6-TG compared to untreated control. Three independent RNASeq experiments were conducted, and average data were plotted using GraphPad Prism V9. **B-H.** Time-kill assays performed using 0.5/1× MIC concentrations of CLOX (2 or 4 µg/ml), 5-FU (16 µg/ml), 5-FUrd (1 µg/ml), Gem (0.25 µg/ml), Mito (0.125 µg/ml), 6-TG (128 µg/ml) and H_2_O_2_ (0.18 µM) alone or in combination with glutathione (GLUT; 10mM) and/or CLOX 2 µg/ml. For the 6-TG killing assay CLOX 4 µg/ml was used because 6-TG antagonises CLOX activity. Panels show killing kinetics for: (B) CLOX ± GLUT; (C) 5-FU ± GLUT/CLOX; (D) 5-FUrd ± GLUT/CLOX; (E) Gem ± GLUT/CLOX; (F) Mito ± GLUT/CLOX; (G) 6-TG ± GLUT/CLOX; (H) H_2_O_2_ ± GLUT/CLOX. Exponential phase cultures were inoculated into MHB 2% NaCl at a starting cell density of approximately 1×10^6^ CFU/ml, with or without antibiotics and GLUT as indicated, and CFUs enumerated after 0, 2, 4, 6, 8, 12, and 24 h. The data presented are the average of three independent experiments plotted using GraphPad Prism V9 and standard deviations are shown.

We recently reported an important role for ROS in MRSA killing by antibiotic/adjuvant combinations that included 5-FU and 5-FUrd (17). Here we extended these experiments to examine the role of ROS in MRSA killing by anticancer drugs alone and in combination with CLOX. The minimum concentration of CLOX with bactericidal activity (as measured by a 3-log decrease in CFU/ml) was found to be 4 µg/ml (Fig. 6B). CLOX (2 µg/ml) achieved a 1-log decrease in CFU/ml after 12 h followed by recovery over 24 h (Fig. 6B), presumably due to the acquisition of potentiator mutations. Consistent with previous studies implicating ROS in β-lactam killing of MRSA (53), glutathione partially reversed the significant increase in bactericidal activity of CLOX 4 µg/ml (Fig. 6B). Next JE2 killing by 5-FU, 5-FUrd, Gem and Mito at MIC levels was measured alone and in combination with 0.5× MIC CLOX (2 µg/ml) (Fig. 5C-F). On their own, none of these drugs achieved significant (3 log) MRSA killing. However, when combined with CLOX (2 µg/ml), significant killing was measured (Fig. 6B-E). Furthermore, this killing was partially reversed by glutathione, also implicating ROS in the mechanism of action of these combinations. In contrast to 5-FU, 5-FUrd, Gem and Mito, time kill experiments with 6-TG again demonstrated its antagonism of CLOX at 4 µg/ml, and no significant change was observed with glutathione (Fig. 6G). As a control the ROS inducer H_2_O_2_ was also shown to increase MRSA killing by CLOX via a mechanism that could be reversed by glutathione (Fig. 6H). Together, these findings demonstrate that ROS generation is a contributor to the bactericidal activity of cloxacillin/anticancer drug combinations against MRSA.

### Drugs targeting nucleotide metabolism inhibit biofilm formation and eradicate established biofilms, with variable effects on hemolytic activity

A dose-dependent inhibition of JE2 biofilm formation was measured using 0.5×, 1×, and 2× MICs of the nucleotide analogue drugs, reaching significance at 2× MIC (Fig. 7A). Biofilm inhibition by Mito and 5-FU at 2× MIC was similar to that achieved by proteinase K which has previously been shown to inhibit and disperse MRSA biofilms (54) (Fig. 7A).

**Fig. 7.**
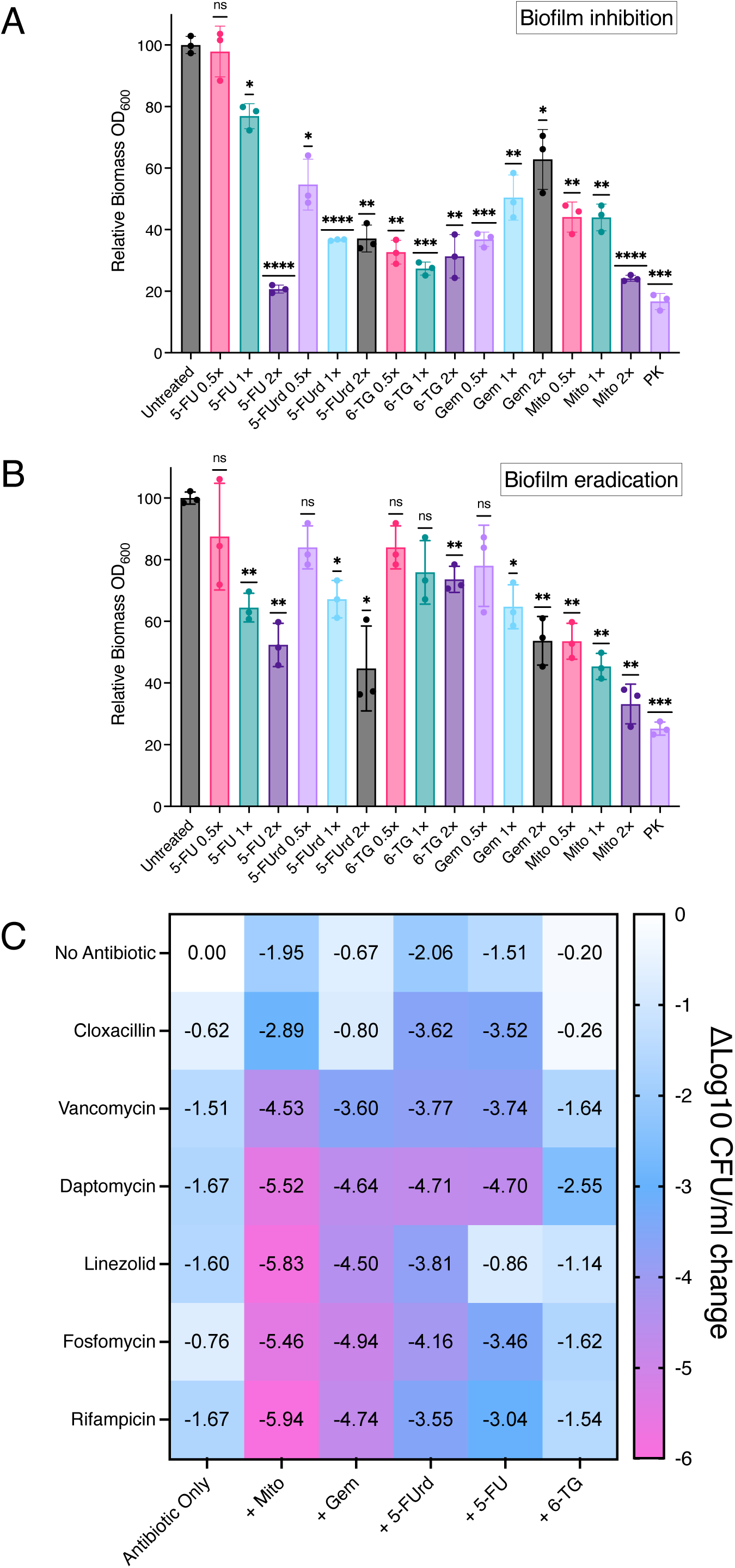
Combinations of antimicrobial and anticancer drugs significantly inhibit and inactivate MRSA biofilms. **A.** Inhibition of JE2 biofilm formation by anti-cancer compounds at 0.5×, 1×, and 2× MICs. Biofilms were grown in BHI 1% glucose for 24 h and biomass quantified by crystal violet staining (OD_600_) and normalized to the untreated control (100%). Data represent means ± SD from ≥3 independent experiments. Statistical significance was determined using GraphPad Prism v9 (P<0.05, *P<0.01, **P<0.001, ***P<0.0001). **B.** Pre-formed 24 h JE2 biofilms were treated with anti-cancer compounds (0.5×, 1×, and 2× MICs) for an additional 24 h before being washed, quantified by crystal violet staining (OD_600_) and normalized to the untreated control (100%). **C.** Pre-formed 24h JE2 biofilms were treated for an additional 24 h with antimicrobial and anticancer drugs singly and in combinations at concentrations approximating reported C_max_ values. Surviving cells were serially diluted, enumerated on MHA, and the results presented as Δlog_10_ CFU/ml relative to untreated controls. The antimicrobial/anticancer drug concentrations used were: cloxacillin (89 µg/ml); vancomycin (50 µg/ml), daptomycin (100 µg/ml), linezolid (27 µg/ml), rifampicin (10 µg/ml), gemcitabine (40 µg/ml), mitomycin C (5 µg/ml), 5-fluorouridine (200 µg/ml), 5-fluorouracil (68 µg/ml) and 6-thioguanine (78 µg/ml). The data, which represent the mean of 3 independent experiments, were plotted using GraphPad Prism v9.

These drugs also exhibited a dose-dependent ability to disperse established JE2 biofilms, with 2× MIC Mito achieving similar levels of eradication to proteinase K (Fig. 7B). This Mito concentration (2 µg/ml) is lower than the C_max_ (5 µg/ml (55)). 6-TG did not disperse biofilms at 0.5× MIC or 1× MIC and although it did reduce biofilm biomass by approximately 25% at 2× MIC (512 µg/ml), this concentration is significantly higher than the C_max_ (78 µg/ml) (56).

Next, we measured inactivation of 24 h JE2 biofilms by combinations of these anti-cancer drugs with daptomycin and rifampicin, which are recommended for the treatment of biofilm infections (57, 58), vancomycin, daptomycin and linezolid, which are used in the treatment of MRSA infections (59, 60), as well as cloxacillin and fosfomycin which act synergistically with 5-FU, 5-FUrd, Mito and Gem. Enumeration of colony forming units after treatment with these combinations increased in the inactivation of JE2 biofilms (as determined by enumeration of colony forming units, CFUs) compared to the antimicrobials or anticancer agents alone (Fig. 7C). Notably, Mito combined with rifampicin, linezolid, daptomycin, fosfomycin or vancomycin achieved >5 log reductions in biofilm viability (Fig. 7C), which meets the threshold for significant antibiofilm activity (61). Combinations of Gem, 5-FU and 5-FUrd with rifampicin, linezolid, daptomycin, fosfomycin or vancomycin achieved 3-4.9 log reductions in biofilm viability (Fig. 7C). Combinations of CLOX with the anticancer drugs did not achieve the levels of biofilm eradication observed for the other antibiotics (Fig. 7C). Taken together these data reveal the potential of anti-cancer/antibacterial drug combinations to significantly inhibit biofilm formation and reduce biofilm viability, which may in turn explain increased dispersal of mature biofilms.

6-TG has previously been shown to exhibit anti-hemolytic activity (30). To assess the effects of 5-FU, 5-FUrd, Gem and Mito on hemolysis, cell-free supernatants from cultures grown for 20 h in the presence of 0.125× MIC of each drug were incubated with whole sheep blood, and hemolytic activity was quantified (Fig. S4). Both 6-TG and Mito strongly reduced hemolytic activity, while 5-FU produced a very modest yet significant decrease. In contrast, 5-FUrd and Gem had no significant effect compared to the MHB control. These findings highlight the differential effects of nucleotide analogue drugs on MRSA hemolytic activity and, importantly, indicate that these agents do not enhance virulence potential.

## Discussion

Nucleotide antimetabolites used as anti-cancer drugs interfere with nucleotide homeostasis and prevent DNA synthesis leading to cell death (62). While some anticancer drugs including 5-FU, 6-TG and Mito have previously been reported to have antibacterial properties (30, 63, 64), their potential has been largely neglected due to concerns about potential toxicity (65). However, deploying nucleotide analogues for the treatment of acute bacterial infections may require shorter treatment durations (66, 67) and/or lower doses, particularly if they exhibit synergy with existing antibiotics. Building on our previous discovery that purine nucleosides and mutations in the purine salvage pathway modulate β-lactam resistance in MRSA (16), the data presented in this study reveal that nucleotide analogue drugs modulate MRSA susceptibility, biofilm formation, and virulence-associated phenotypes.

The pyrimidine-targeting agents 5-FU, 5-FUrd, Gem, and Mito potentiated the activity of β-lactams and fosfomycin, whereas the purine analogue 6-TG antagonized the activity of these antibiotics against MRSA. Consistent with these phenotypes, 5-FU, 5-FUrd, Gem, and Mito downregulated genes involved in lysine, glutamate, and arginine biosynthesis, whereas 6-TG induced their expression. This opposing response aligns with the antagonistic effect of 6-TG relative to the purine nucleosides guanosine and xanthosine, which potentiate β-lactam activity (16). Notably, lysine biosynthetic genes were among the most differentially expressed following treatment with nucleotide analogue drugs. Given that lysine and glutamate are key components of the *S. aureus* peptidoglycan stem peptide (68), their repression is consistent with the observed synergy with β-lactams, whereas their activation by 6-TG supports its antagonistic effect. The strong repression of autolytic activity by 6-TG, in contrast to the minimal effects of pyrimidine-targeting agents, further supported mechanistic differences between these compounds. Several studies have highlighted the relationship between autolytic activity and β-lactam susceptibility, although the precise mechanistic impact of these phenotypes on each other remains unclear (69). Repression of autolysis by 6-TG and increased β-lactam resistance is consistent with the model proposed by Fisher and Mobashery in which the bactericidal activity of β-lactams is dependent on deregulated Atl activity at the cell division septum (70).

Transcriptomic analysis showed that the anticancer drugs tested have physiological effects beyond DNA damage. Overall, the expression of ≥133 genes were significantly regulated ≥2-fold after exposure to the anti-cancer compounds, including regulators of DNA damage repair and oxidative stress responses. The pleiotropic effects of the pyrimidine antimetabolites also correlated with the ability of 5-FU, 5-FUrd and Mito to potentiate the activity of fosfomycin and the ability of Mito to synergise with daptomycin. Furthermore the predicted impact of these drugs on DNA damage and induction of ROS was associated with altered regulation of DNA damage response genes and the ability of the ROS scavenger glutathione to reverse the anti-MRSA activity of 5-FU, 5-FUrd, Gem and Mito on their own and in combinations with OX. In contrast, 6-TG did not significantly alter expression of DNA damage response genes, and its anti-MRSA activity was unaffected by glutathione.

The nucleotide analogue drugs also possess intrinsic antibiofilm activity against MRSA and can potentiate the efficacy of clinically relevant antibiotics. The potent inhibition of biofilm formation and dispersal of established biofilms by Mito and 5-FU, at concentrations within or below clinically achievable levels, suggests that disruption of nucleotide metabolism may target biofilm-specific processes. The enhanced killing observed in combination with rifampicin, daptomycin, linezolid, fosfomycin, and vancomycin, used in the treatment of MRSA infections (57–60), indicates that these agents sensitize biofilm-associated cells to antibiotic-mediated killing, potentially by increasing metabolic vulnerability or altering cell envelope homeostasis. Combinations using therapeutically relevant concentrations of antibiotics and anti-cancer drugs were able to achieve 5-6 log reduction in biofilm viability, which was comparable to previously reported anti-biofilm therapeutics (61). In contrast, the limited activity of 6-TG, particularly at clinically relevant concentrations, further highlights mechanistic differences among nucleotide analogues.

Finally, despite their broad physiological effects, the nucleotide analogue drugs did not enhance MRSA virulence-associated phenotypes. Consistent with previous reports showing that 6-TG reduces Hla expression (30) and improves necrotic lesion healing in murine models (44), we also observed that 6-TG repressed hemolytic activity in MRSA culture supernatants. Mito also reduced repressed hemolytic activity, whereas hemolytic activity was largely unaffected by 5-FU, 5-FUrd and Gem.

Together, these findings demonstrate that pyrimidine-targeting nucleotide analogues potentiate anti-MRSA antibiotics via modulation of cell wall biosynthesis, oxidative stress, and biofilm survival, supporting their potential repurposing as adjunctive therapies against difficult-to-treat MRSA infections.

## Materials and Methods

### Bacterial strains, antibiotic minimum inhibitory concentration (MIC) and synergy assays

The *S. aureus* strains used in this study, JE2 (71), MW2 (72), BH1CC (73) and COL (74) were routinely grown in Mueller-Hinton Broth (MHB), Mueller-Hinton Agar (MHA), Tryptone Soy Broth (TSB) or Tryptic Soy Agar (TSA) (all Oxoid). MIC measurements were conducted according to CLSI guidelines (75). The medium was supplemented with 50 mg/l calcium chloride for all experiments measuring daptomycin susceptibility in accordance with Clinical and Laboratory Standards Institute (CLSI) guidelines (76, 77) and 25 mg/l glucose-6-phosphate for fosfomycin susceptibility assays (78). Strains were grown for 24 h on MHA and 5-6 colonies were resuspended in 0.85% saline and the cell density adjusted to OD_600_ = 0.1. This cell suspension was diluted 1:20 in 0.85% saline from which 10 µl used to inoculate 100 µl of MHB supplemented with serially diluted antibiotics. Microdilution checkerboard assays to measure synergy between 5-FU, 5-FUrd, 6-TG, Gem, Mito and oxacillin or cloxacillin were performed in 96 well plates as described previously (16, 79, 80).

### Growth and drug susceptibility assays using a Tecan microplate reader

A Tecan Sunrise microplate reader was used to record growth (OD_600_) every 15 minutes for 24 h using Magellan software. *S. aureus* strains were streaked on MHA, grown for 24 h before 5-6 colonies were suspended in 0.85% NaCl. This cell suspension was adjusted to OD_600_ = 0.1 from which 20 µl was inoculated into wells containing 180 µl of MHB or MHB containing anticancer compounds corresponding to 0.5×, 1× and 2× MICs of the corresponding strain. For Acy, 5-FC and AZT which exhibit MIC values ≥1024 µg/ml, 1024 µg/ml was used in these experiments.

### Isolation and genome sequencing of MRSA mutants with increased drug resistance

50 µl aliquots from overnight cultures of JE2 were used to inoculate 5 ml MHB containing 0.5× MICs of 5-FU, 5-FUrd, 6-TG, Gem, Mito, which were then grown for 24 h. These cultures were serially passaged every 24 h in MHB supplemented with 0.5× MICs of the anticancer drugs until turbid growth was observed. Single colonies from the turbid cultures were isolated on MHA.

### Whole genome sequencing

Genomic DNA (gDNA) extractions were carried out as described previously (59). Whole genome sequencing was performed commercially by Novogene, using Illumina short read technology. CLC Genomics Workbench software (Qiagen Version 20) was used for genome sequencing analysis. A reference genome was prepared by mapping wild-type JE2 reads to the closely related USA300 FPR3757 genome sequence (RefSeq accession number NC_07793.1). Read sequences from mutants with increased resistance to 5-FU, 5-FUrd, Gem, Mito and 6-TG were mapped onto the assembled JE2 sequence and single nucleotide polymorphisms (SNPs), deletions or insertions were identified as performed previously (59).

### Preparation of RNA samples for transcriptomic analysis

Total RNA was extracted for transcriptional analysis as described previously with minor adjustments (83). JE2 overnight cultures were diluted 1:100 in 10 ml fresh MHB and allowed grow to exponential phase (OD_600_ = 0.4). 1× MICs of 5-FU, 5-FUrd, Gem, Mito and 6-TG were added to the cell suspensions, which were then incubated for a further hour before the cells were pelleted at 7,000 ×g for 5 min at 4 °C. Next the cell pellets were resuspended in 1 ml RNAlater (Sigma) and incubated at ambient temperature (∼18 °C) for RNA preservation. The remaining steps were carried out on ice. The cells were again pelleted at 7,000 ×g and resuspended in 1 ml RNA lysis buffer before being transferred to lysis matrix B tubes for mechanical disruption (40 s at 6.5 m/s, thrice) in a MP Biomedical FastPrep bead beater instrument. RNA was extracted using a Qiagen RNeasy mini-kit according to the manufacturer’s instructions. Isolated RNA samples were treated with Turbo DNase (Invitrogen) according to manufacturer’s instructions. Sample quality and quantity were evaluated using Nanodrop, Qubit and agarose gel electrophoresis analysis. The purified RNA samples were sequenced commercially by Novogene who also performed the subsequent bioinformatic analysis. rRNA was removed using Illumina Ribo-Zero Plus rRNA Depletion Kit before proceeding to cDNA synthesis. Libraries were pooled and sequenced on an Illumina NovaSeq6000 instrument using a strategy based on paired-end reads with 150bp reads (PE150). Raw reads were mapped to reference genome USA300 FPR3757, log2 fold changes were calculated and plotted using GraphPad Prism V9.

### Lysostaphin susceptibility assays

JE2 overnight cultures were inoculated 1:100 into fresh 25 ml MHB 2% NaCl supplemented with 0.125 × MICs of oxacillin (8 µg/ml), MICs of 5-FU (4 µg/ml), 5-Furd (0.25 µg/ml), Gem (0.0625 µg/ml), Mito (0.03 µg/ml), 6-TG (32 µg/ml) as indicated and grown for 3 h. Cells were centrifuged 1 min 12,000 ×g and resuspended in TSM buffer (0.05 M Tris HCl pH7.5, 10 mM MgCl2, 0.5 M sucrose) to a cell density of approximately 1×10^8^ CFU/ml. TSM suspensions with or without lysostaphin were incubated for 90 mins, and serially diluted to enumerate colony forming units (CFU/ml) on TSA plates.

### Autolytic activity assays

Samples (250 µl) from overnight cultures were inoculated into 25 ml of MHB, grown at 37°C (200 rpm) to an OD_600_ of 0.5, centrifuged at 4°C, washed with 20 ml of cold PBS, resuspended in 5 ml of cold PBS, and then adjusted to an OD_600_ of 1. Then, 200 µl of the cell suspension was transferred to a 96 well plate, and Triton X-100 was added at a final concentration of 0.1% (vol/vol). The initial OD_600_ was recorded before incubation at 30°C with shaking (200 rpm). Thereafter, the OD_600_ was recorded every 15 min for 4 h using Tecan Sunrise Plate reader at 30°C, and autolytic activity is expressed as a percentage of the initial OD_600_. Three biological replicates were performed for each condition.

### Checkerboard/Synergy assays

Ninety-six well plates were utilized for checkerboard assays, one compound was diluted vertically while the other compound tested was diluted horizontally, resulting in 100 µl of media per well. Strains were streaked onto MHA plates and grown for 24 h at 37 °C. Five-ten colonies were resuspended into 0.85% NaCl and adjusted to 0.5 McFarland standard (OD_600_ = 0.1). The cell suspensions were then diluted 1:20 in 0.85% NaCl and 10 µl used to inoculate 100 µl media. Plates were grown for 24 h at 37 °C, OD_600_ values were then measured using Tecan Sunrise microplate reader.

### Fosfomycin susceptibility assays

JE2 overnight cultures were inoculated 1:100 into fresh 25 ml MHB 2% NaCl supplemented with 0.125× MICs of oxacillin (8 µg/ml), MICs of 5-FU (4 µg/ml), 5-Furd (0.25 µg/ml), Gem (0.0625 µg/ml), Mito (0.03 µg/ml), 6-TG (32 µg/ml) as indicated and grown for 3 h. Cells were centrifuged 1 min 12,000 ×g and resuspended in PBS, samples were washed once more with cold PBS. Cells were adjusted to a cell density of approximately 1-5 × 10^8^ CFU/ml in 5ml TSB supplemented with fosfomycin and grown at 37°C (200 rpm). CFUs were enumerated on TSA plates after 0,1, 2, 3, 4, 5, 6 h.

### Fluorescent confocal microscopy analysis using HADA and SYTO 9

Overnight JE2 cultures grown in MHB were diluted 1:100 in 5 ml fresh MHB and grown for 4h in subinhibitory (1/64 MIC) of OX, 0.125× MICs of 5-FU, 5-FUrd, Gem, Mito or 6-TG or a combination of OX and these drugs before being incubated with HADA (500 µM final concentration) in the dark for 5 mins. Cells were then pelleted for 2 mins at 14,000 ×g and resuspended in PBS supplemented SYTO 9 (1 µM final concentration) at cell density OD600 = 1.0 before being incubated for 15 mins at 37 °C (in the dark). Cells were washed twice with 1 ml PBS before being resuspended in 1 ml PBS (OD600 = 1.0) and 5 µl aliquots of these cell suspensions were spotted onto a thin 1% agarose patch prepared in PBS and the stained bacteria imaged at 1000× magnification using an Olympus LS FLUOVIEW Fv3000 Confocal Laser Scanning Microscope. Representative images of 3 independent biological samples were analysed using Fiji (ImageJ). HADA and SYTO 9 fluorescence intensities of individual cells were quantified using Fiji (ImageJ) for 90 cells per condition (30 cells per biological replicate). Septal division defects were recorded of dividing cells were quantified using Fiji (ImageJ) for 180 cells per condition (60 cells per biological replicate).

### MRSA killing assays

Killing assays were performed according to CLSI guidelines (84), as previously described (16). For cloxacillin killing assays, the overnight cultures were grown in MHB 2% NaCl, diluted 1:100 and grown for 3 h before being inoculated into 25 ml of MHB 2% NaCl alone or MHB 2% NaCl supplemented with 0.25-0.5× MICs of CLOX (2 or 4 µg/ml), 0.5× MICs of 5-FU (16 µg/ml), 5-Furd (1 µg/ml), Gem (0.25 µg/ml), Mito (0.125 µg/ml), 6-TG (128 µg/ml), or H_2_O_2_ (0.18 µM) alone or in combination with glutathione (10mM). CFUs were enumerated on TSA plates after 0, 2, 4, 6, 8, 12, and 24 h.

### Biofilm inhibition and dispersal assays

Overnight cultures were diluted 1:200 into BHI supplemented with 1% glucose and 200 µl aliquoted into the individual wells of a 96-well Nunclon Delta-treated, flat-bottom microplate. For inhibition assays 5-FU, 5-FUrd, Gem, Mito or 6-TG were added at 0.5×, 1× and 2× MICs and the plates incubated for 24 h at 37 °C. For dispersal assays, biofilms were first grown at 37 °C for 24 h. Next the spent media was removed and replaced with fresh media supplemented with 5-FU, 5-FUrd, Gem, Mito or 6-TG at 0.5×, 1× and 2× MICs, before being incubated for a further 24 h at 37 °C. Finally, the remaining biofilm was carefully washed twice with 125 µl PBS and the biofilms fixed by drying at 65 °C for 1 h before being stained with 200 µl 0.4% crystal violet for 15 mins. After removal of excess crystal violet by dunk washing in water, the plates were air dried upside down on tissue paper before the crystal violet in the remaining biofilms was extracted in 30% acetic acid. The absorbance of the crystal violet solutions extracted from the biofilms was measured spectrophotometrically at 600 nm in a Tecan Sunrise microplate reader to indirectly quantify the remaining biomass compared to the untreated control. Proteinase K (2 µg/ml) was used as a positive control to inhibit biofilm formation and to disperse MRSA biofilms (85–87).

### Biofilm eradication assays

Biofilm eradication assays were performed as described previously (16, 88). Briefly, biofilms were grown in 2 ml BHI 1% glucose inoculated into the wells of 24-well tissue culture coated plates at 37℃ for 24 h before being carefully washed twice with PBS. BHI supplemented with antibiotics as indicated was then added to the biofilm wells and the biofilms incubated for a further 24 h at 37℃. The biofilms were dispersed by scraping, serially diluted and CFUs enumerated on TSA plates. The antibiotic concentrations used in these experiments was equal to or less than the C_max_ in humans as follows: 89 µg/ml cloxacillin (89), 50 µg/ml vancomycin (90), 100 µg/ml daptomycin (91), 27 µg/ml linezolid (92), 300 µg/ml fosfomycin (89), 10 µg/ml rifampicin (93), 40 µg/ml gemcitabine (94), 5 µg/ml mitomycin C (55), 200 µg/ml 5-fluorouridine (95), 78 µg/ml 6-thioguanine (56), 68 µg/ml 5-fluorouracil (96).

### Quantitative sheep blood hemolysis assay

Quantitative sheep blood hemolysis assays were performed as described previously (82). Overnight JE2 cultures were diluted 1:100 into fresh MHB supplemented with subinhibitory concentrations (0.25× MICs) of 5-FU, 5-FUrd, Gem, Mito and 6-TG and/or oxacillin (1/64 MIC) for 20 h at 37 °C shaking at 200 rpm. The cells were pelleted at 5,500 ×g for 5 mins and 100 µl of supernatant was added to 25 µl whole sheep blood which had been added to 875 µl PBS, and the mixture incubated at 37 °C for 30 mins. Following centrifugation at 5,500 ×g for 1min, the supernatant was removed, transferred to spectrophotometer cuvettes and the absorbance OD_543_ measured. 100 µl of 1% Triton and 100 µl of PBS were used as positive and negative controls, respectively.

## Author Contributions

Aaron Nolan: Conceptualization, Formal analysis, Funding acquisition, Investigation, Methodology, Writing - original draft, Writing - review & editing.

Sarah Byrne: Methodology, Writing - review & editing.

Merve S. Zeden: Conceptualization, Formal analysis, Funding acquisition, Project administration, Investigation, Data curation, Methodology, Supervision, Writing - original draft, Writing - review & editing, Project management.

James P. O’Gara: Conceptualization, Formal analysis, Funding acquisition, Project administration, Supervision, Writing - original draft, Writing - review & editing, Project management.

## Acknowledgements.

This research was supported by grants from Health Research Board (ILP-POR-2019-102), Research Ireland (19/FFP/6441) and the Joint Programming Initiative for Antimicrobial Resistance/Health Research Board Ireland (<u>PURIFY AMR</u>) to J.P.O’G, a Thomas Crawford Hayes research award (2023) to A.C.N., a Research Ireland (SFI-IRC) Pathway Fellowship (22/PATH-S/10804) to M.S.Z., SDG Research Seed Funding from the University of Galway College of Science and Engineering to M.S.Z and a Research Ireland Government of Ireland Postgraduate Scholarship (GOIPG/2025/8574) to S.B. We are grateful to Dr Peter Owens from the Centre for Microscopy & Imaging facility for assistance with confocal microscopy. The funders had no role in study design, data collection and interpretation, or the decision to submit the work for publication.

---

**Table S1.**
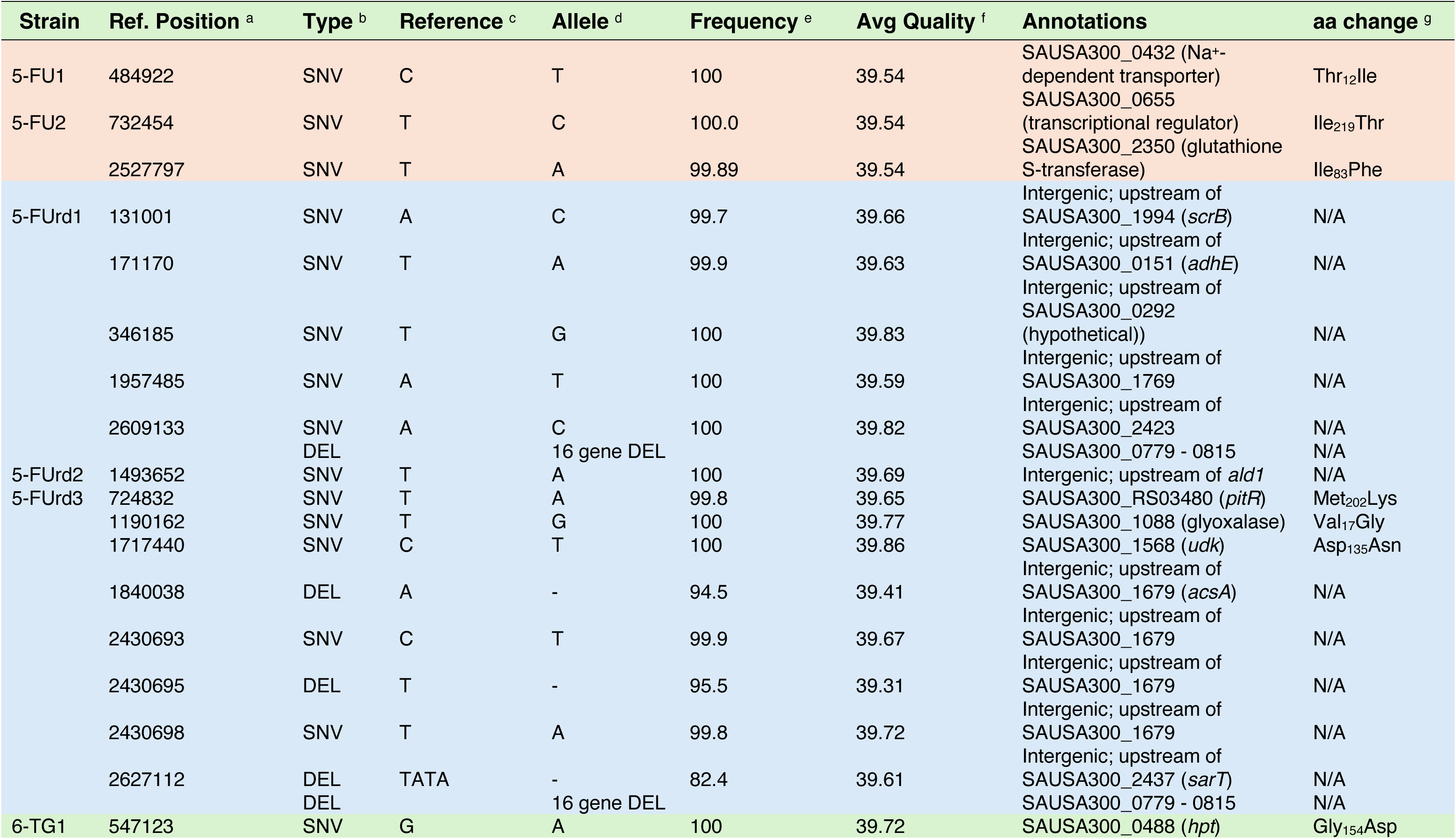
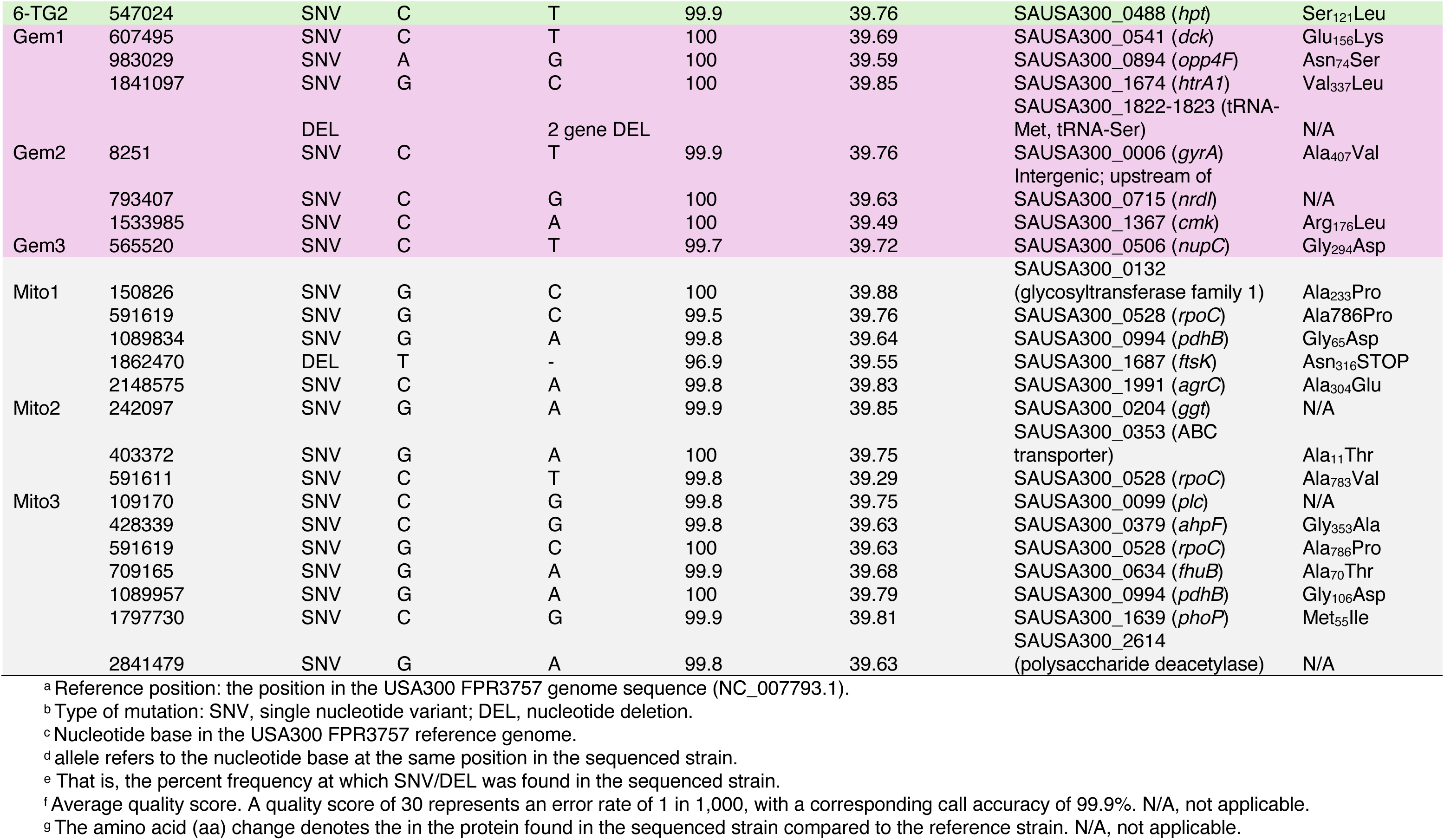
Genomic changes in mutants of MRSA strain JE2 with increased resistance to 5-FU, 5-FUrd, 6-TG, Gem and Mito.

